# Genomic V-gene repertoire in reptiles

**DOI:** 10.1101/002618

**Authors:** David N. Olivieri, Bernardo von Haeften, Christian Sánchez-Espinel, Jose Faro, Francisco Gambón-Deza

## Abstract

Reptiles and mammals diverged over 300 million years ago, creating two parallel evolutionary lineages amongst terrestrial vertebrates. In reptiles, two main evolutionary lines emerged, one gave rise to Squamata, while the other gave rise to Testudines, Crocodylia and birds. In this study, we determined the genomic variable (V)-gene repertoire in reptiles corresponding to the three main immunoglobulin (Ig) loci and the four main T-cell receptor (TCR) loci. We show that squamata lack the TCR *γ/δ* genes and snakes lack the V*κ* genes. In representative species of testudines and crocodiles, the seven major Ig and TCR loci are maintained. As in mammals, genes of the Ig loci can be grouped into well-defined clans through a multi-species phylogenetic analysis. We show that the reptile VH and V*λ* genes are distributed amongst the established mammalian clans, while their V*κ* genes are found within a single clan, nearly exclusive from the mammalian sequences. The reptile and mammal V-genes of the TRA locus cluster into six common evolutionary clans. In contrast, the reptile V-genes from the TRB locus cluster into three clans, which have few mammalian members. In this locus, the V-gene sequences from mammals appear to have undergone different evolutionary diversification processes that occurred outside these shared reptile clans.

## 1. Introduction

The order Reptilia is believed to have evolved from amphibian-like creatures, the labyrinthodonts, which date back to the Carboniferous period some 312 million years ago (Mya) (Laurin & Reisz, 1995; Laurin & Girondot, 1999). The descendants of those precursors became progressively more terrestrial and independent of their aquatic origins, venturing out on a larger expanse of land with many new different habitats (Benton & Donoghue, 2007; deBraga & Rieppel, 1997; van Tuinen & Hadly, 2004; Reisz & Müller, 2004). Evolving from this primordial amphibious ancestor, a lineage began that would later split into two separate lines, the Sauropsids and the Synapsids, the latter of which would eventually give rise to the line of extant mammals (Benton, 2005; Hedges & Kumar, 2009). Shortly after this separation of the Sauropsids and Synapsids, the sauropsids further divided into two separate lineages. One gave rise to the present day Squamata (lizards and snakes) and Tuatara, while the other gave rise in a very early period to Testudines, later to Crocodylia, and finally, gave way to birds (Benton, 2005; Hedges & Kumar, 2009).

The first jawed vertebrates developed an adaptive immune system consisting of B and T lymphocytes together with their associated antigen receptors, immunoglobulins (Ig) and T-cell receptors (TCR), respectively. This adaptive immune system evolved in parallel within Reptilia and the sister group of mammals from their common amphibian ancestor (Janes et al., 2010).

Although Igs and TCRs share a basic modular structure (Charlemagne et al., 1998; Schroeder & Cavacini, 2010; Narciso et al., 2011), Igs are made up of four polypeptide chains (two identical larger or heavy (H) chains and two identical smaller or light (L) chains), while TCRs are heterodimers (Schroeder & Cavacini, 2010). The H chains of Igs consist typically of four or more globular domains, with the N-terminal domain varying amongst the Igs of different B-cell clones (termed variable (V) domain), and the remaining domains being shared by many B-cell clones (termed constant (C) domains). In contrast, L chains of Igs and TCR polypeptide chains consist of two domains, an N-terminal V domain, and a C domain. In addition, there are two different kinds of TCRs, one composed of *α/β* heterodimers and the other of *γ/δ* heterodimers. In both Igs and TCRs, only the V domains interact with antigen.

Each chain of the Igs and of TCRs is encoded in different loci. Jawed vertebrates have three main loci corresponding to Ig genes (IGH, coding for H chains, and IGK and IGL, coding for *κ* and *λ* L chains, respectively), and four main loci associated with the TCR genes (TRA, TRB, TRG and TRD, coding for TCR *α*, *β*, *γ* and *δ* chains, respectively) (Cannon et al., 2004; Danilova & Amemiya, 2009; Flajnik & Kasahara, 2010; Sun et al., 2012). In all these loci, multiple unique V gene sequences (V-genes) exist that share a common homology. Each Ig and TCR V domain is encoded by a unique combination of a V-gene, a D segment (only in Ig H and TCR *β* and *δ* chains) and a J segment, the largest contribution to the V domain amino acid sequence being from the V-gene. Thus, identifying the V-gene repertoire within the genome of a species provides information about its potential for mounting an adaptive immune response against antigen.

The Ig and TCR loci may have undergone an independent evolution, because they are located in different chromosomes, or in widely separated regions within the same chromosome. One exception to this general trend exists, the TRD locus, which is embedded within the TRA locus. This independent evolution hypothesis is supported by the formation of V-gene clusters in phylogenetic trees constructed from these sequences (Olivieri et al., 2013). If true, this explanation would imply the existence of an ancestral gene (or genes) that define each of the extant Ig and TCR V-genes.

In reptiles, published studies of the Ig genes, together with the functionality of their products, are scarce (Gambón-Deza et al., 2012; Magadán-Mompó et al., 2013; Magadán-Mompó et al., 2013). At present, there are no studies of V-genes in the the TCR loci of reptiles. In Squamata, all the species studied so far have IgM, an IgD consisting of eleven domains, and IgY (Sun et al., 2011; Gambón-Deza et al., 2012). In Anole, the V*κ* and V*λ* chains have been described and the absence of the IGK locus in snakes (Wei et al., 2009; Gambón-Deza et al., 2009) has been confirmed. Testudines have a more complex IGH locus structure (Li et al., 2012; Xu et al., 2009; Magadán-Mompó et al., 2013), having one IgM gene, one IgD gene encoding eleven constant domains, and multiple genes for IgY and IgD2. Previous studies of the IGH locus in crocodiles have described three genes for IgM, one gene for IgD encoding seven viable constant domains, three IgA genes, three IgY genes that are similar to IgY in birds, and two IgD2 genes, which encode chimeric Igs with IgA and IgD related constant domains (Magadán-Mompó et al., 2013). These studies have also detected in crocodiles that the IgA heavy chain is similar to the IgX and IgA of amphibians and birds, respectively.

In contrast to the growing literature related to Igs in reptiles, there are fewer studies of TCRs in these species. In this article, we uncover the genomic V-gene repertoire of both immunologic receptors in species representative of the three main orders of extant Reptilia: Squamata, Testudines, and Crocodylia, and show relations that provide a deeper understanding of the evolution of this large vertebrate lineage (Modesto & Anderson, 2004; Benton, 2005).

## 2. Methods

For these studies, we used genome data from whole genome shotgun (WGS) assemblies that are publicly available at the NCBI and provided in FASTA format in the form of contigs. Until now, complete V-gene annotations have only been available in two species (i.e., human and mouse), while incomplete annotations have existed in a few species, and even in these cases, only within certain loci. We used our VgenExtractor software tool (Olivieri et al., 2013) to obtain the V-gene sequences from genome files. Our software algorithm searches and extracts the genome sequences of functional V-genes based upon known motifs. In particular, the algorithm scans large genome files and extracts candidate exon sequences that fulfill specific criteria: they are flanked by recombination signal sequences, they have a valid reading frame without stop codon, the length is at least 280 bp long, and they contain two canonical cysteines and a tryptophan at specific positions.

From the set of translated candidate exons produced by the VgenExtractor tool, a fraction of these sequences could pass the initial filter, purely by chance, but have structures that are quite different from a functional V-gene. Such sequences are easily discarded by subjecting all the candidate sequences to a BLASTP comparison against a V-gene consensus with a modest threshold (e.g., evalue = 10^−15^). This operation produces a set of exons possessing the structural requirements for being functional V-genes.

Once viable V-genes are obtained we need to classify them into their respective loci. The corresponding amino acid sequences were analyzed with the suite of tools provided by Galaxy (Goecks et al., 2010). The analysis workflow of the amino acid sequences consist of a multiple BLASTP alignment against a set of consensus sequences, each representing one of the seven main Ig and TCR V-gene loci. Each candidate V-gene sequence is assigned to the locus having the highest score with respect to the corresponding consensus.

We developed another independent algorithm for classifying V-genes. In this method, given a V-gene sequence, we determine to which locus this sequence most likely belongs by obtaining a heuristic score calculated from the results of a BLASTP comparison against the NR protein database. The score is a weighted combination of information from sequence similarity results (or hits). In particular, the score is constructed from the following: (a) the relevance and content words from the protein description, (b) the similarity score, c) and the sequence coverage. A score is calculated for each locus, and the V-gene sequence is assigned to the most likely locus. This method has been validated from known annotations in the human and mouse (Olivieri et al., 2013). This technique is used to show the consistency of results obtained with the the simpler workflow described above and can be used to resolve ambiguous loci predictions of the simpler method, which may arise in rare cases.

To compare the repertoire of V-genes in extant species, we carried out single and multi-species phylogenetic analysis. The sequences for this study are available at a new public database repository vgenerepertoire.org (Olivieri & Gambón-Deza, 2014). At present, approximately 20,000 V-gene sequences from 82 mammals and 12 reptiles are indexed. For both the single and multispecies phylogenetic analysis, sequences were aligned using Clustal-omega (Sievers & Higgins, 2014) and trees were constructed with FastTree (Price et al., 2010) (using maximum likelihood and WAG matrix). We used MEGA5 (Tamura et al., 2011) for visualization.

In *Crocodylidae* we used the NCBI accession numbers for the following species: American alligator (*Alligator mississippiensis*; AKHW000000001), Chinese alligator (*Alligator sinensis*; AVPB0000000001), while the following species used assemblies found in www.crocgenomes.org: saltwater crocodile (*Crocodylus porosus*; croc sub2.assembly), and gharial (*Gavialis gangeticus*; PRJNA172383). In *Testudines* we used the NCBI accession numbers for the following species: spiny softshell turtle (*Apalone spinifera*; APJP00000000.1), green sea turtle (*Chelonia mydas*; AJIM00000000.1), Chinese soft-shelled turtle (*Pelodiscus sinensis*; AGCU00000000.1), painted turtle (*Chrysemys picta bellii*; AHGY00000000.1). In *Squamata* we used the NCBI accession numbers for the following species: green anole (*Anolis carolinensis*; AAWZ00000000.2), Burmese python (*Python bivittatus*; AEQU000000000.2), the king cobra (*Ophiophagus hannah*; AZIM00000000.1), and data from the Boa Assemblathon 2 (Bradnam & et.al., 2013) (*Boa constrictor*; scaffold 6C file).

## 3. Results

From the WGS data obtained from the NCBI repositories of twelve Reptile species, we used the *VgenExtractor* tool to obtain functional V-genes and classify them with respect to the locus to which they belong. Table 1 presents a summary of all the functional V-genes encountered in the analysis. For many of these species, it is the first time that Ig and TCR V-genes have been described, and it completes the sparse data presently available with respect to these genes.

**Table 1:**
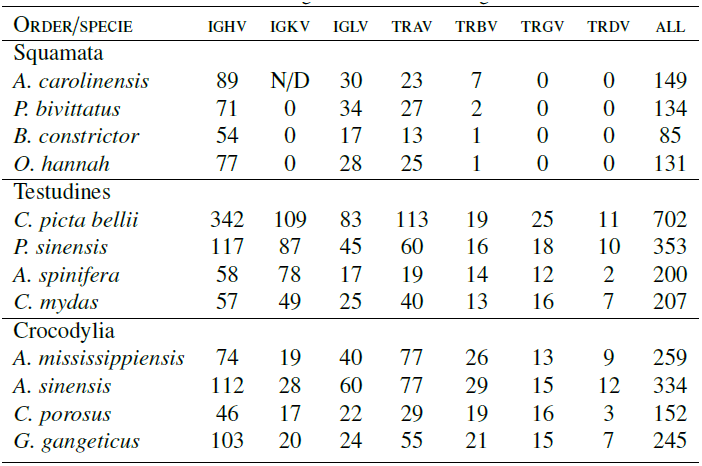
Distribution of V-genes across the main Ig and TCR loci

### 3.1 Squamata

Within the Squamata order, we studied the genomes of one Iguanidae (i.e., *Anolis carolinensis*), and three snakes (i.e., *Python bivittatus*, *Ophiophagus hannah* and *Boa constrictor*). The phylogenetic trees of the V-genes for these species are shown in Figure 1.

**Figure 1:**
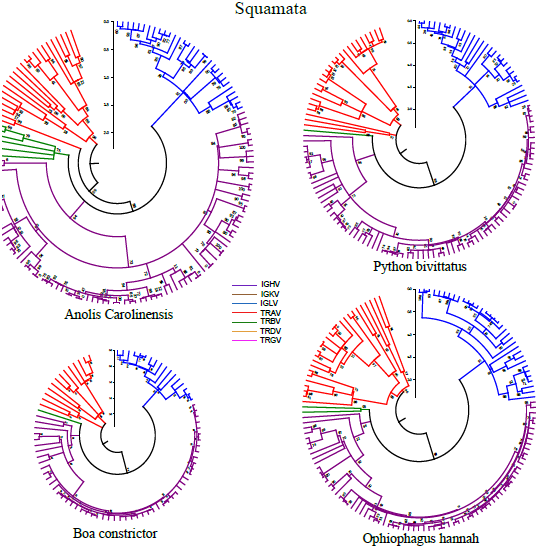
V-gene repertoire in Squamata. Trees were constructed with the amino acid sequences obtained from the VgenExtractor tool. These sequences were aligned with Clustal-omega. For constructing the phylogenetic trees, a maximum likelihood algorithm with the WAG matrix and 500 bootstrap replicates were realized for validation. Rooting was performed at the midpoint and the linearization provided by Mega5 was applied to improve the visualization of the trees. For each sequence, two independent classification techniques (view Material and Methods) were used to determine its respective locus, thereby confirming that clades of the tree correspond to independent loci of Ig and TCR.

Within the antibody V genes of the Anole, we did not detect the V*κ* genes, contradicting previous studies (Alfoldi, 2011; Wei et al., 2009). A further analysis showed that we did not detect these V*κ* sequences because they lack tryptophan at position 41 (Figure 2). This result is surprising, since the presence and position of this amino acid is a general characteristic of all functional V domains described so far, implying a structural alteration in this chain. The snake species, which arose later in the evolution of this lineage, no longer possess the *κ* chain genes. A possible explanation for this loss in snakes may be due to this structural modification of the *κ* sequence found in older Squamata. For the case of the IGH locus in the Anole, the results agree with those we described previously by an independent method (Gambón-Deza et al., 2009).

**Figure 2:**
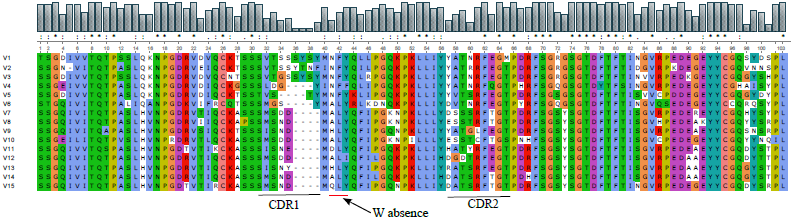
Alignment of amino acid sequences deduced from V-genes obtained manually in the genome of *A. carolinensis*. As can be seen, the amino acid tryptophan (W) does not occupy the canonical position 41, representing an anomaly as compared with other functional V-genes. We produced the alignment and graphical output with the *Seaview* software program (Gouy et al., 2010).

With respect to TCR genes, our analysis showed that the Anole possesses a low number of V genes in the TRB locus. Also, we found that V genes corresponding to the *γ* and *δ* chains (TRGV and TRDV genes) are absent. Because of this anomalous finding, we searched the Anole genome for genes that encode the constant (C) regions of the TCR *γ* and *δ* chains, with negative results. We also analyzed transcriptome sequences for this species obtained from the liver and kidney (available from the NCBI public repository: Anole liver: SRA064777, trace SRR648821, Anole kidney: SRA059267, trace SRR579557). While there were many Ig and TCR *α*/*β* mRNA sequences found in these organs, we obtained negative results when searching for TCR *γ*/*δ* sequences. It is possible that cells having TCR *γ*/*δ* may be abundant in other specialized tissues where transcriptome data is lacking. Nonetheless, our studies, carried out with data from different tissue and different sequencing methods, produced null results for searches of V-genes or constant gene sequences for TCR *γ* and *δ* chains. Thus, these results strongly suggest that these species have lost the TCR *γ*/*δ* receptor.

In the three serpents studied, there are few V-genes (Table 1). Most of these sequences from these species belong to the IGH and IGL loci (Figure 1). In the three snakes, we found that the IGK locus does not exist. From a phylogenetic analysis, we found that V sequences from the IGH locus belong to a single group or clan that has undergone a recent expansion. This result is different from that found in *A. carolinensis*, where many V-gene clans are found in the IGH locus. In the case of the IGL locus, the number of V-genes in Serpentes is similar to that in the Anole. Finally, just as with *A. carolinensis*, no V-gene sequences corresponding to the TRG and TRD loci were found and only one or two V genes were identified in the snake TRB locus.

From studies we carried out previously in mammals, we found a higher birth and death rate amongst the V-genes of the Ig loci as compared to that in the TCR loci. Trees of V-gene sequences in reptiles have the same structure, suggesting that the same mechanisms exist. Figure 3 shows recent duplications that occurred within the IGH and TRA loci. These loci are contained within chromosome segments of length 610 Kb and 140 Kb, respectively. In this figure, track duplications are drawn in green and represent those sequences having a similarity greater than 90 percent for segment sizes > 1Kb. As can be seen, there are many duplications in the IGH locus and these duplication processes occurs specifically in the IGHV region and not in the constant (i.e., IGHC) region. By applying the same similarity criteria, few duplication tracks are visible in the TRAV region, and those that are visible have smaller segment sizes: ≤ 0.3 Kb. Finally, we can observe that some duplications coincide with V-genes (indicated with red lines) in the IGHV region, while no V-genes coincide with the duplication tracks shown in the TRAV region.

**Figure 3:**
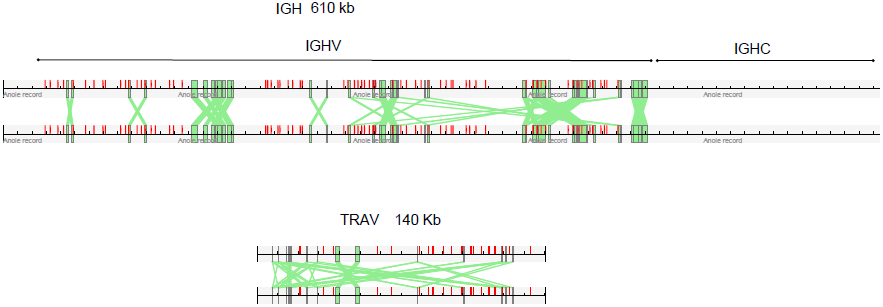
Recent duplications in IGHV (top) and TRAV (bottom) loci in the Anole. Track duplications tracks are drawn in green. Constraints for duplications in: (a) the IGH locus are: segments ≥ 1Kb, identity ≥ 90%; (b) the TRA locus are: segments ≤ 0.3Kb; identity ≥ 90%. Duplications only exist in segments where V-genes are located (not in constant regions), and in the case of the IGH locus, some coincide with the position of V-genes, indicated by the red lines.

### 3.2. Testudines and Crocodylia

In the differentiation of the reptile lineage to birds, the Testudine and Crocodylia branches emerged. Four Testudine genomes are available: those of three land tortoises, *Chrysemys picta bellii*, *Pelodiscus sinensis* and *Apalone spinifera*, and one marine turtle, *Chelonia mydas*. Figure 4 shows the phylogenetic analysis of the V genes sequences from these species. For the western painted turtle, *C. picta bellii*, we found more than 700 V-genes, being the specie with the most V-genes that we have studied to date. By comparison, the marine turtle has ≈ 1/3 as many genes (Table 1). In all four turtles, there is a preponderance of Ig V-genes as compared with the number of TCR V-genes. In the three Ig loci, there is an expansion of multiple V-gene clans, similar to what happens in the Anole, suggesting that the structure of these genes is more diverse than that found in mammals (Gambón-Deza et al., 2009). Within the Ig loci, there are more Ig heavy chain genes than those for light chains; the *λ* light chain has the least Ig V-genes. With respect to TCR V-genes, the TRA locus have the largest set, while the TRB locus has on average much less genes. Such a low amount of V*β* genes is observed in all reptiles, opening the possibility that an underlying process exists. In contrast to the Squamata, each of the turtle species we studied possess TCR *γ* and *δ* V-genes, albeit with a wide variance amongst them. From the phylogenetic study of these V-genes, the TRGV genes are included in one specific clan, while the TRDV genes are within the clan of the TRAV genes (Figure 4).

**Figure 4:**
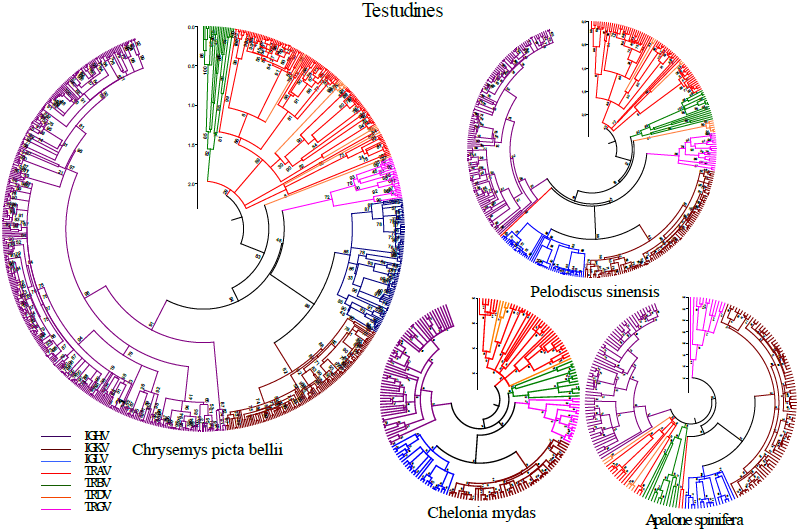
V-genes in Testudines. The trees were constructed in the same way as described in Figure 1.

In the order Crocodylia, there are four publicly available genomes (*Alligator mississippiensis*, *Alligator sinensis Crocodylus porosus*, and *Gavialis gangeticus*). In each species, we found V-regions of the three main Ig loci and four TCR loci (Table 1). There are more V-genes for Ig than for TCR. Amongst the Ig V-genes, the majority of them are within the IGH locus. All four Crocodylia species have higher numbers of V-genes in the IGL locus than in the IGK one. Figure 5 shows the phylogenetic trees of the V-genes for each of these species.

**Figure 5:**
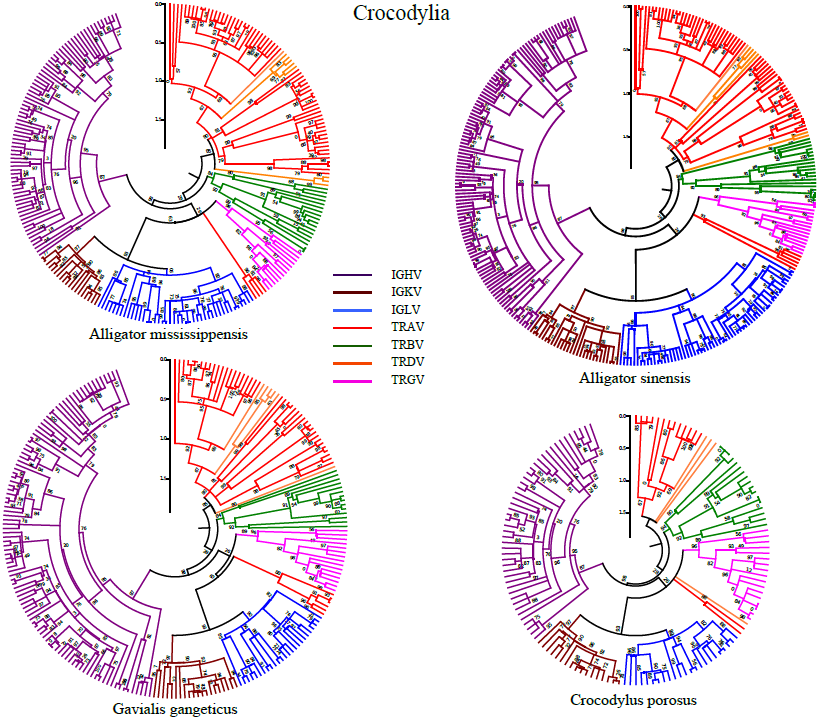
V-genes in Crocodylia. The trees were constructed in the same way as described in Figure 1.

### 3.3. Clans in V-gene sequences

We studied the phylogenetic relationships of V-genes from the 82 mammal and 12 reptile species available in Vgenerepertoire.org (Olivieri & Gambón-Deza, 2014). From the large set of immunoglobulin V gene sequences in this study, there is evidence for distinct groups or clans (Lefranc & Lefranc, 2001) of related sequences from different species amongst the clades. Previous studies have identified such clans in human and mice; the IMGT (Lefranc et al., 2009; Giudicelli & Lefranc, 2012) has annotated three clans amongst the VH genes, three in V*κ*, and at least five clans in V*λ* (Lefranc & Lefranc, 2001; Giudicelli & Lefranc, 1999).

Figure 6 shows multi-species trees obtained from alignments of the V-genes from the Ig loci. Since the purpose of these trees was to explore the clan structure, we used all available sequences of reptiles and mammals instead of trying to retain the same number of species from each class. In these plots, the sequences corresponding to mammals are represented in blue, while those for reptiles are drawn in red. The nodes having sequences which belong to clans annotated by the IMGT are labelled in the trees. As seen in the trees of Figure 6, many sequences do not belong to a previously defined clan. Since the clan studies to date have been based upon a limited set of species, the set of identified clans has been necessarily inadequate. It is only now, with the availability of large number of V-genes sequences from a broad representation of mammal and reptile species (Olivieri & Gambón-Deza, 2014), that it is possible to define new clans.

**Figure 6:**
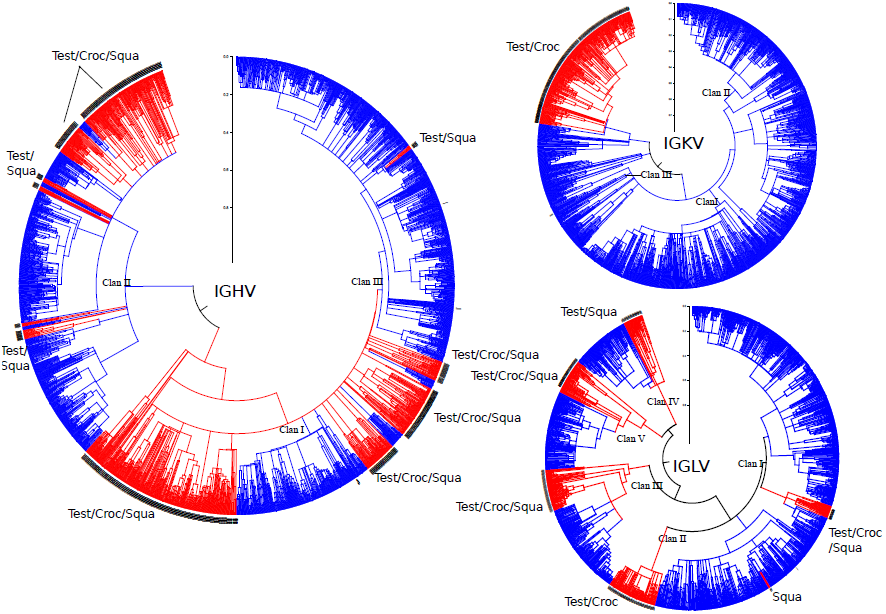
Trees were generated from 11 reptiles and 50 mammals using the V-gene sequences of the three Ig loci. The clans with reptile taxa are marked in red while those for mammals are indicated in blue. Previous studies (Lefranc & Lefranc, 2001; Giudicelli & Lefranc, 1999) have defined the presence of the clans indicated in the plots. The alignment of the amino acid sequences was performed with clustal-omega and tree was constructed using FastTree with the WAG matrix. Plots were constructed using MEGA v5.2 with the linearized branch option. The labels of the Reptilia orders correspond to Testudines (*Test*), Crocodylia (*Croc*), and Squamata (*Squa*).

In the IGHV tree of Figure 6, the V-gene sequences of reptiles are grouped together and these groups are interspersed within mammal clans. For example, Clan-II contains sequences from Testudines, Crocodylia, and Squamata that together form subgroups, and these subgroups are interposed with several mammalian subgroups comprising many sequences. The tree for the IGLV sequences shows that the reptile V*λ* sequences are also grouped in several subgroups and these are distributed amongst the five mammalian clans previously defined by the IMGT. In the IGK locus, more V*κ* clans can be identified in mammals than previously described, so that the present clan nomenclature is not sufficient to group all the taxa. All the reptile IGKV genes cluster into a single group forming an independent clan. Also, this clan contains only a small group of mammal V-gene sequences, indicating a different evolution in this locus between mammals and reptiles.

V-gene clans have not been previously identified within the TCR loci. The multi-species trees of Figure 7, consisting of TCR sequences, demonstrate that the clan concept is also relevant amongst these loci. Indeed, in the TRA locus, six clans can be identified, each having sequences from both reptiles and mammals. As can be seen in the figure, the roots of each of the clans correspond to reptile sequences, indicating the maintenance of these sequences over longer periods of time than those found in mammals. The clans are numbered according to the amount of taxons in the clan, ranked from largest (Clan I) to smallest (Clan V). The tree also demonstrates that clans I and II diversified significantly in mammals, while in others, such as clan V, there was less diversification in mammals than witnessed in reptiles. Also, we can deduce the presence of additional clans in reptiles that retain remnants of sequences that were later lost in mammals.

**Figure 7:**
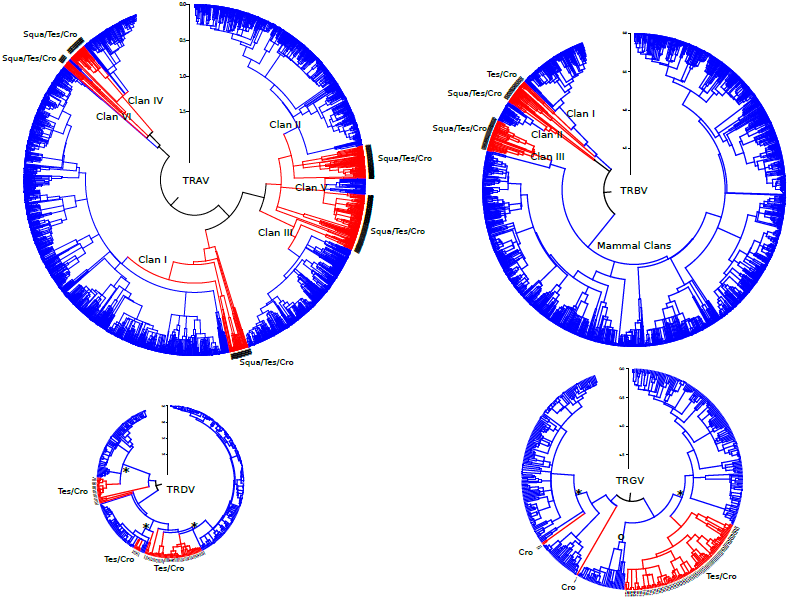
Trees were generated from 11 reptiles and 50 mammals using the V-gene sequences of the four TCR loci. The clans in reptile taxa are marked in red while those for mammals are in blue. The nodes that root the mammal clans with the reptile clans are marked by (*), while others that are exclusively from mammals or reptiles are marked with (o). The alignment and tree construction was performed in a similar way to that of Figure 6. The labels of the Reptilia orders correspond to Testudines (*Test*), Crocodylia (*Croc*), and Squamata (*Squa*).

In the tree for the TRB locus in Figure 7, three distinct clans can be identified that group mammal and reptile taxons. Nonetheless, the majority of mammalian sequences in this locus do not share the same clan with those from reptiles. Few mammal sequences are found within the reptile clans, while those grouped in separate clans appear to have undergone an evolutionary diversification different from reptiles within this locus.

Trees from the TCR *γ*/*δ* have a similar structure. In the TRDV tree, three clans can be identified that contain sequences from mammals, Testudines, and Crocodylia (this locus does not exist in Squamata, as described previously). Similar results are seen in the multi-species tree for the TRG locus, but in this case, the majority of sequences from reptiles belong to a single clan.

## 4. Discussion

How did the genes that code for Igs and TCRs arise and evolve in lineages of reptiles? It is known that extant reptiles and mammals are separate vertebrate lineages that descended from a common ancestor. These vertebrates must have inherited their antigen recognition machinery, namely Ig and TCR genes, from this common ancestor. These vertebrates evolved on land in different habitats, and underwent further speciation. After the divergence from mammals, the reptile lineage split in two. One of these lines led to present day Squamata, while the other gave way to Testudines and Crocodylia. We found that the Ig V-gene repertoire in reptiles is similar in structure to that found in mammals. Just as in mammals, the data suggests that Ig V-genes underwent a diversification process with high birth/death rates.

Another striking result is that the Squamata species lost the TCR *γ*/*δ* locus. Neither V-nor C-gene sequences were found in either the genome or several transcriptome datasets, indicating the absence of these genes. In each of the Squamata species we studied, there is a low number of V-genes in both the TRA and TRB loci, perhaps underscoring a particular characteristic of their TCR *α*/*β*. This study also revealed several relevant facts with regard to the evolution of the Ig loci. From *A. carolinensis*, we found that the V*κ* sequences have an anomalous alteration in the canonical tryptophan, whose presence is a general characteristic of all functional V-genes described so far. This alteration observed in Iguanidae may be related to the loss of this locus in the suborder of Serpentes. It is known that birds also lost the IGK locus. However, in the evolutionary line that gave rise to Testudines and Crocodylia, common to that of birds, we did not detect this loss of tryptophan 41 at the canonical position. It is thus surprising that two evolutionary lines, serpents and birds, that diverged ∼275 Mya independently lost the IGK locus.

In the Testudines and Crocodylia lineage, there is no such gene loss as seen in the Squamata. Table 1, shows the V-gene distribution across all the seven major Ig and TCR loci. While the average number of TRB V-genes is larger than that found in Squamata, it is still less than the average number found in mammals (Olivieri & Gambón-Deza, 2014). For the case of the TRG and TRD loci, the average number of V-genes was similar to that found in mammals.

At the evolutionary level, there are studies that show that the VH genes are grouped into three clans (Lefranc & Lefranc, 2001; Giudicelli & Lefranc, 1999; Lefranc et al., 2009). These studies distributed the sequences in clans from limited information based upon a small number of mammal species. In this work, we presented multi-species trees formed from the V-genes of more than 90 mammal and reptile species. These trees show that additional clans exist beyond those described from previous annotations.

In a previous analysis of *A. carolinensis* (Gambón-Deza et al., 2009), we showed that its VH gene sequences cluster into five distinct groups, one of which is quite different from the rest. The necessity to define new clans is shown in the reptilian VH and V*κ* sequences. In the V*κ* genes of Testudines and Crocodylia, there exists a single clan different from those of mammals and within which mammals have only a few V-gene sequences (Figure 6). This suggests that reptiles generated *κ* chains from a homogeneous ancestral group, from which mammals either diversified very little or lost most of the variants that were generated. Both in serpents and in birds, the *κ* chain was lost, indicating that the V*κ* genes expressed in reptiles may be dispensable. In contrast, the IGKV tree of Figure 6 demonstrates the large *κ* chain diversification that took place in mammals, corresponding to the most common Ig chain in these species, such as the case in human and mouse.

The retention of the five clans in the V*λ* gene sequences, as described previously in mammals, is significant because it suggests a positive selection mechanism. The reptile sequences maintain this distribution of five V-genes clans found in mammals and each is rooted with a reptile clan. The difference in sequences (which may be structural) that conditioned the grouping within clans has been maintained in both lineages for more than 300 Mya, indicating functional transcendency not yet understood.

Instead of recognizing antigen directly, TCRs recognize a complex of antigen-derived peptide and a major histocompatibility complex (MHC) molecule. Thus, this restriction could be accounted for by a co-evolution selection between the TCR V-genes with MHC molecules. At present, there are few comprehensive studies exist that characterize the MHC molecules in reptiles (Glaberman et al., 2008; Glaberman & Caccone, 2008; Glaberman et al., 2009).

In a previous publication (Olivieri et al., 2013), we commented on the presence of long branches in the trees corresponding to the TCR V-gene sequences of mammals. This same phenomena is also observed in the TCR V-gene trees for reptiles. Due to the ability to study a large number of V-gene sequences in these loci, we also describe for the first time the presence of well-defined clans in the multi-species trees. In the TRA locus, the reptiles and mammals share common clans and the roots of the clades are from both reptiles and mammals, indicating a common origin. One explanation for this observed phenomena is a structural or functional preservation of products from this loci in species that diverged more than 300 Mya.

The TRBV multi-species tree of Figure 7 describes an evolution between reptiles and mammals that is quite different from that seen in the TRAV tree. The V*α* sequences evolved from common ancestors, as seen by the shared mammal and reptile root nodes. The distribution of the V*α* sequences are also evenly distributed amongst the clans. This same homogeneous distribution is not seen in the TRBV tree. In particular, the vast majority of mammal V*β* sequences have diversified into separate clans (i.e., mammal clans). In this way, the V*β* sequences of reptiles group within tree old clusters with little representation from mammals. Thus, contrary to V*α* sequences, reptile and mammal V*β* sequences are segregated from each other, in a way reminiscent of V*κ* sequences. While the common evolutionary features of V*α* in reptiles and mammals supports a coevolution hypothesis between TCR and MHC, the evolutionary differences observed in the V*β* suggests this chain may have a different role in reptiles and mammals. Studies of MHC evolution in reptiles could provide clues that clarify this issue.

